# Robust, single-shot, optical autofocus system utilizing cylindrical lenses to provide high precision and long range of operation

**DOI:** 10.1101/2022.07.14.498938

**Authors:** J. Lightley, S. Kumar, E. Garcia, Y. Alexandrov, C. Dunsby, M.A.A. Neil, P.M.W. French

## Abstract

We present a robust, “real-time” optical autofocus system for microscopy that provides high accuracy (<230 nm) and long range (∼130 µm) with a 1.4 numerical aperture oil immersion objective lens. This autofocus can operate in a closed loop, single-shot functionality over a range of ±37.5 µm and can also operate as a 2-step process up to ±68 µm. A real-time autofocus capability is useful for experiments with long image data acquisition times, including single molecule localization microscopy, that may be impacted by defocusing resulting from drift of components, e.g., due to changes in temperature or mechanical drift. It is also vital for automated slide scanning or multiwell plate imaging where the sample may not be in the same horizontal plane for every field of view during the image data acquisition. To realise high precision and long range, we implement orthogonal optical readouts using cylindrical lenses. We demonstrate the performance of this new optical autofocus system with automated multiwell plate imaging and single molecule localisation microscopy and illustrate the benefit of using a superluminescent diode as the autofocus light source.

## 1. Introduction

Thermal and other fluctuations can lead to drift of microscope components over time such that the desired field of view (FOV) becomes defocused. This is a particular problem for microscopy experiments with long image data acquisition times, including single molecule localization microscopy (SMLM), and is an essential consideration for automated slide scanning or multiwell plate imaging where the sample may not be in the same horizontal plane for every field of view during the image data acquisition. The precision of an autofocus system should be better than the depth of field of the microscope. When using high numerical aperture (NA) objective lenses, this precision should be less than ∼500 nm.

Autofocus functionality can be realised using software-based autofocus techniques, e.g. ^1^, that analyse images acquired in the microscope in the normal way and calculate the degree of defocus (i.e. the distance of the desired sample plane from the imaging plane (usually the focal plane of the objective lens). Typically, this entails acquiring an image of the sample and using previously recorded image data from an axial scan of the sample or a test object to determine the defocus. Often this requires the acquisition of new image data at each new FOV, which can increase phototoxicity and slow the imaging throughput. The range of this autofocus approach typically scales with the confocal parameter of the objective lens and rarely exceeds 10 µm. This can be a problem when moving between FOV in multiwell plates^2^ or slide-scanning.

For more challenging applications such as slide-scanning and high content analysis, a hardware-based autofocus or “optical autofocus” is often implemented. This entails incorporating an additional optical system in the microscope that typically tracks the motion of the microscope coverslip/sample interface. Defocus may be measured through secondary imaging of features in the sample^3^ or fiduciary markers^4,5^ (such as gold beads) but more often is measured using an autofocus laser beam focused on the coverslip/sample interface with the back reflected light either being monitored using a photodiode detector or being imaged onto a secondary “autofocus camera”. The use of low power near infrared radiation for the autofocus can significantly reduce the phototoxic light dose on the sample compared to software-based techniques using primary microscope images. Defocus can be quantified using a metric calculated from an image of this back-reflected laser beam on a secondary (autofocus) camera (such as displacement of images^6^ or centroid^7^). Such calculated metrics typically do not distinguish whether the defocus is positive or negative. This may be addressed by offsetting the focusing of the back-reflected laser beam at the autofocus camera such that any drift of the sample relative to the microscope objective does not cause the reflected laser beam to pass through focus (minimum spot size) on the autofocus camera^8^. Unfortunately, this decreases the operating range of the optical autofocus.

The precision and operating range of the autofocus can be adjusted by controlling the diameter of the autofocus laser beam in the back aperture of the microscope objective lens – and therefore the autofocus confocal parameter. Normally this leads to a compromise between precision and range of the optical autofocus system. In a recent paper^2^, however, we relaxed this compromise by using a slit to shape the autofocus laser beam such that it exhibited a different confocal parameter in the planes parallel and perpendicular to the slit. This enabled us to realise an accuracy better than the depth of field of the objective lens (∼600 nm) over an operating range of ∼+/-100 µm. We further used a convolutional neural network (CNN) to determine the defocus by comparing the autofocus camera image to training data of autofocus camera z-stacks previously recorded when scanning a coverslip through the focal plane of the microscope objective lens. This CNN-based readout was sufficiently sensitive to the variation of the autofocus camera images that it could also determine the sign of defocus. However, we found that, because the optical system of the autofocus itself was subject to drift, it would only work well with training data acquired within a few hours. We overcame this by collecting separate training data over multiple days and forming a composited training data set that encompassed the perturbed variations in the alignment of the autofocus system (and therefore the images recorded on the autofocus camera.

Using this composite training data, the CNN-based optical autofocus has functioned for >12 months. However, it is a significant effort to set up the system and acquire the composite training data and it may be necessary to acquire new training data if the microscope objective lens was changed. We therefore decided to develop a new optical autofocus that retains the capability for high precision and long operating range but does not require machine learning and the associated acquisition of training data. It also senses the sign of defocus without compromising the operating range and can operate in closed loop for real-time focus control.

## 2. Methods

Figure 1 shows the first configuration of this new optical autofocus system^9^, which was implemented on an *openFrame*^10^ microscope configured for easySTORM^11^ using a 100x objective lens of 1.4 NA (Olympus UplanSApo 100x 1.40 oil). It is similar to our previously published configuration^2^ except that the autofocus laser beam is being collimated after emerging from a single mode optical fibre by a pair of cylindrical lenses of different focal length orientated at 90° to each other to produce the desired elliptical collimated beam (instead of using a spherical collimating lens and a slit^2^). By using cylindrical lenses with different focal lengths, we can adjust the autofocus confocal parameter in orthogonal directions and thus the precision and operating range of the autofocus. For subsequent readouts of defocus, we binned the autofocus camera images in the directions parallel to the two cylindrical lens axes to generate respective intensity projections.

**Figure 1.**
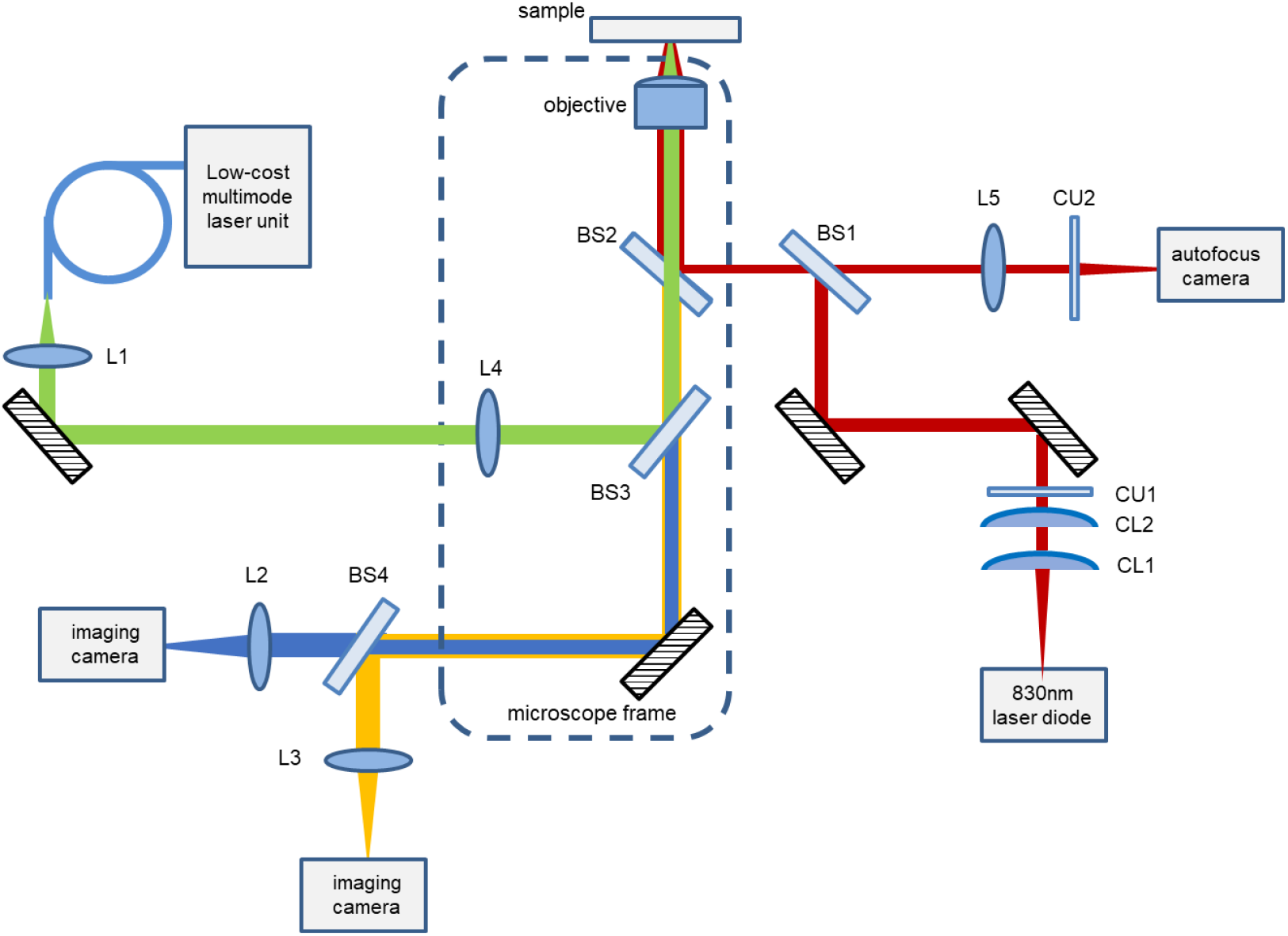
Implementation of the cylindrical lens autofocus on the openFrame. A superluminescent laser diode (Superlum S-930 nm), was coupled into a single mode fibre with a 0.1 NA and the output of the fibre was used as the source for the autofocus. The beam was collimated in one of the orthogonal axes with a 6.4mm focal length cylindrical lens and in the other direction a 20.0mm focal length cylindrical lens to produce an elliptical beam profile. The beam was reflected first by a 50:50 beamsplitter, BS1, and then reflected by a 800 nm short pass filter, BS2. The rectangular beam was then focused onto the sample by an Olympus 100x objective 1.4 NA and reflected at the refractive index change between the coverslip and the sample’s media. The reflected signal then propagated back through the system and was transmitted by the 50:50 beamsplitter, BS1, before being imaged by an auxiliary autofocus camera, (FLIR Chameleon3 CM3-U3-31S4M-CS) by a 200mm focal length lens L5.

The defocus can be determined from a single autofocus camera image by previously recording a z-stack of autofocus camera images as the sample coverslip is translated axially through focus and calculating the FWHM of the Fourier transform power spectrum of each projection at each z-position (defocus value) to create a look-up table of this metric as a function of defocus. We chose to use this image metric to quantify defocus as we found it to be less sensitive to background noise, e.g., from spurious back reflections of autofocus laser light from other components in the optical system, than the FWHM or standard deviation of the autofocus camera intensity projections. In general, we avoid image metrics based on displacement of the back-reflected autofocus laser beam because these can be more sensitive to changes in the optical alignment.

We can directly determine the sign of defocus by placing one of the cylindrical lenses at slightly more or less than its focal length from the optical fibre, such that the peak of the defocus image metric is shifted as a function of defocus. Figure 2 shows a plot of the defocus image metric as a function of defocus (axial displacement of objective lens relative to the sample coverslip) plotted in red for the (background-subtracted) autofocus camera projections in the plane of the larger collimated beam diameter (shorter autofocus confocal parameter) and in blue for the autofocus camera projections corresponding to the longer autofocus confocal parameter. Also plotted in green is the average intensity of the whole autofocus camera image after subtraction of a background image (recorded with no sample coverslip present). This provides a means to remove some unwanted signals, e.g., arising from back reflections from components in the optical system.

**Figure 2.**
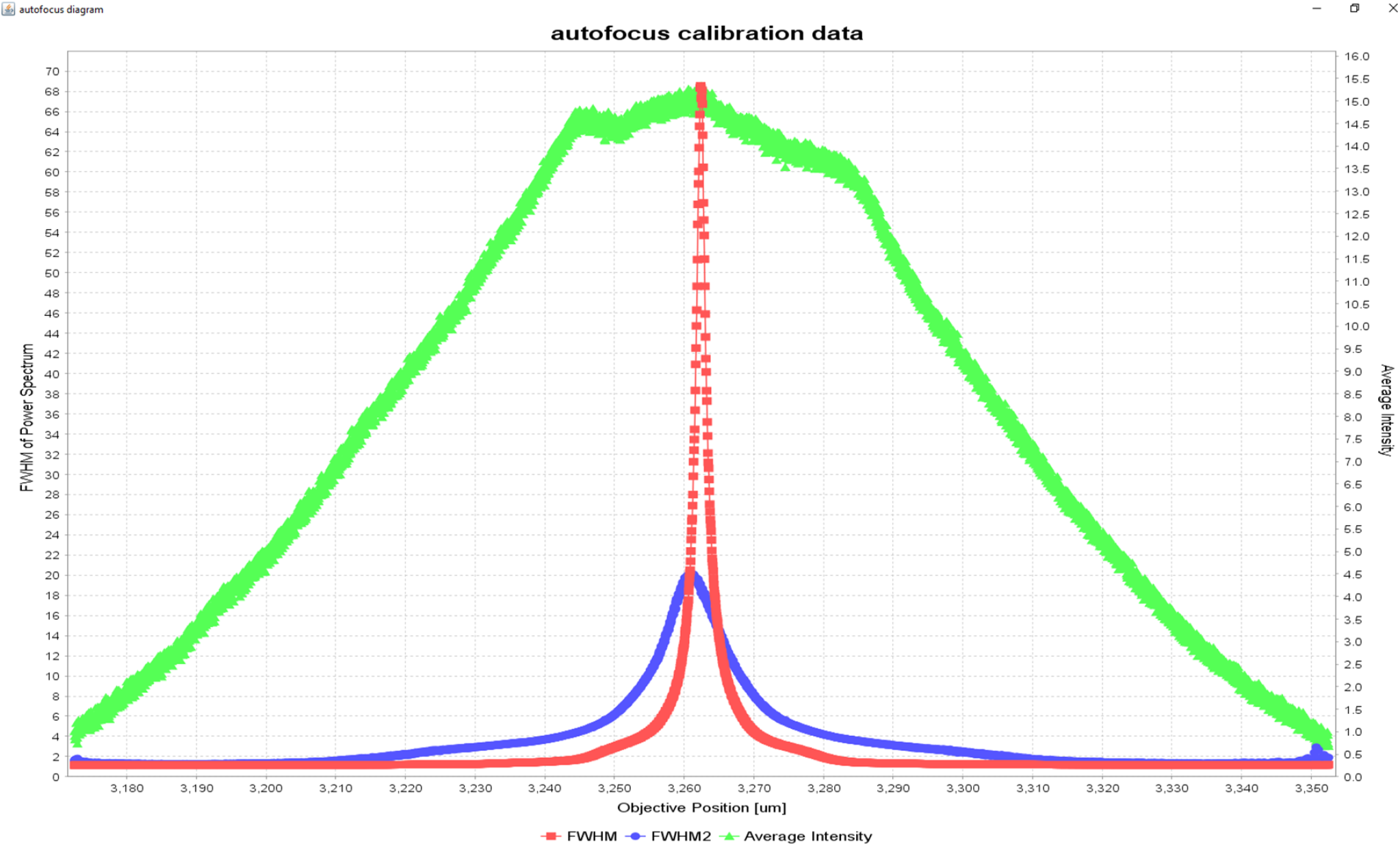
FWHM of the Fourier transform power spectrum of the two projections of average intensity along the horizontal and vertical axes of the sensor of the FLIR Chameleon3 CMOS camera. The red and blue peaks represent the autofocus metrics for the short confocal parameter and long confocal parameter respectively. The green plot shows the average intensity in the image after a background subtraction. As the 20.0mm focal length lens has been positioned slightly further away than its focal length from the fibre, it focuses onto the sample 2 microns axially above the blue peak. This enables closed loop single-shot autofocus

For our previous implementation of an autofocus using a CNN to process the autofocus camera images, the contributions from any unwanted back reflections off components in the autofocus laser path were included in the training image data and did not appear to compromise the system performance. Since this new optical autofocus system is intended to operate without the need for CNN-based analysis and the requisite training image data, these unwanted back-reflections do impact the performance. This is because the high spatial and temporal coherence of autofocus laser beam leads to interference patterns between the main signal reflected from the sample coverslip and any unwanted reflections from other surfaces in the optical system. These interference patterns cause spatial modulation of the image recorded on the autofocus camera and change as the distance from the objective lens to coverslip is varied, e.g., due to drift or an axial scan. Because this unwanted background contribution to the autofocus camera image varies with defocus, it cannot be easily subtracted. To mitigate this, we replaced the single mode diode laser used in our previous autofocus system^2^ with a superluminescent diode (SLD, Superlum S-930 nm) that provides an autofocus laser beam that is spatially coherent but has a short (∼7 µm) temporal coherence length. Thus, only background signals that are back reflected from components within half the coherence length of the coverslip will produce an interference pattern at the autofocus camera and any back reflections from optical components in the autofocus system or from the other side of the coverslip will only contribute to a fixed background, which can be subtracted from the autofocus camera images acquired.

The software to control this optical autofocus system was written as a plugin for *µ-Manager* 2.0. This plug-in enables the user to define the desired in-focus plane and to activate or disable continuous closed loop autofocus correction. Limits can be set to prevent the axial translation of the objective lens or sample beyond pre-determined safe positions if the autofocus system were to fail. A calibration procedure can be run to create the look-up table to associate the autofocus metric values (i.e., FWHM of the Fourier transform power spectrum of the intensity projections) with their respective defocus offset values. A background image can be stored to be subtracted from the autofocus camera image if required. Threshold autofocus metric values can be set to define the regions in which the short range (single-shot) loop autofocus can be used and the regions where the longer range (and multistep) autofocus correction should be used.

## 3. Results

The range over which the autofocus operates is limited by the signal to noise ratio (S/N) of the (red and blue) defocus metric plots. Where the higher precision (red) curve provides sufficient S/N, the autofocus system can operate in single-shot mode, providing rapid focus lock if required. Outside this range, the less precise autofocus can operate over a longer range but can no longer directly determine the sign of defocus. Instead, the system can acquire two autofocus camera images with the relative sample-objective lens position being changed by a few microns between them and then determine the sign and magnitude of defocus. Although less precise, this long range autofocus can move the system back into the range of the more precise (single-shot) autofocus operation.

To determine the operating ranges of the short and long range autofocus functions, the performance of this cylindrical lens-based autofocus system was quantified by automatically imaging 100 nm TetraSpeck fluorescent beads (TetraSpeck, T727) arrayed in a 96-well plate in which each well was coated with poly-L-lysine. The wells were traversed in a snake-like pattern, moving row by row and acquiring a microscope fluorescence image z-stack for one FOV in each well over an axial range of ±4 µm with axial sampling of 100 nm. At each new well, the change in defocus was sufficiently small that the single-shot, high precision autofocus could be first applied to return the microscope to focus and then the fluorescence image z-stacks were acquired. If the autofocus system performed perfectly, then the central image in the fluorescence image z-stack should be at the same axial coordinate as the initial in-focus image for that FOV. To quantify this performance, the fluorescence z-stack image data were analysed using PSFj^12^, previously published software that considers all the individual 100 nm beads appearing in the FOV and determines the slice in the z stack for which each specific bead is in focus. For each FOV, a histogram of the values of the offset of the bead in-focus plane relative to the central plane in the z-stack (i.e., the predicted in-focus plane) can be generated and quantified by the mean and standard deviation of this distribution.

Figure 3(a) presents the performance of the autofocus in each well as a colour map of the defocus error (i.e., the offset between the autofocus-predicted central plane of the z-stack and the actual in-focus plane as determined by PSFj for each bead) and also displays the mean offset of the beads for each FOV. The colour scale indicates the continuous variation of this mean error from 0 µm (green) up to ±2 µm (red). Figure 3(b) presents a histogram of these mean errors/FOV, for which the standard deviation for the 96-well plate is 230 nm and mean of the absolute prediction errors is 179 nm.

**Figure 3.**
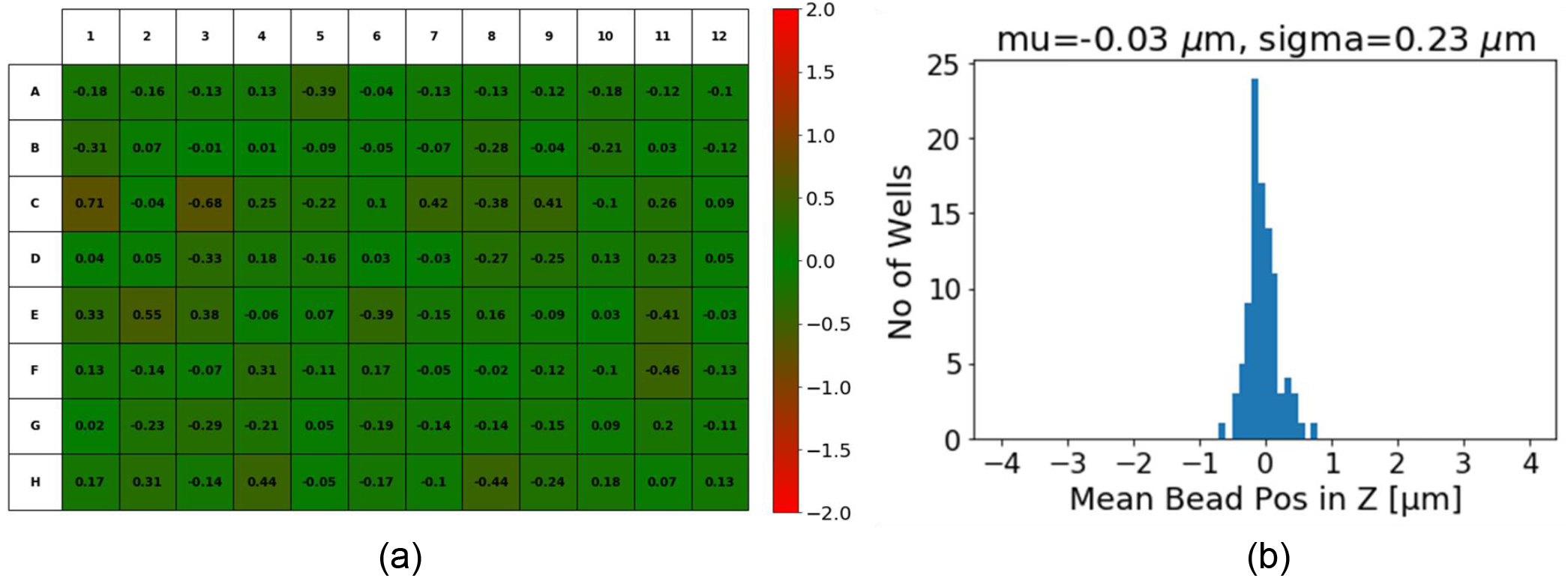
(a) shows a 96 well plate arrayed with 100nm TetraSpeck beads was scanned using autofocus with z-stack acquired in each well. PSFJ was used to determine the in-focus axial plane for each bead and the mean offset from the centre image provided the error in the defocus for each well - shown with associated heatmap of defocus in microns. The false-colour scale shows the mean offset in microns. (b) shows a histogram of errors in defocus for the 96-wellplate.

To further quantify the performance of this cylindrical lens-based autofocus approach, the accuracy of defocus correction from predetermined defocus values was measured. This was done using the reflected image of the autofocus beam from a single FOV and studying the ability of the autofocus system to return the microscope to focus starting from different amounts of defocus. From each pre-set defocus position, the autofocus was applied 10 times and the objective lens was moved to the predicted in-focus position for each of the 10 repeats. To test the shorter range (smaller confocal parameter) autofocus mode, initial defocus values ranging between ±20 µm of defocus in axial steps of 2 µm were used. To test the longer range (larger confocal parameter) mode, initial defocus values ranging between ±60 µm in 10 µm steps were used. These “focus-corrected” objective lens axial positions were compared to the true in-focus objective lens position was determined by acquiring an autofocus camera z-stack and interpolating between frames to find the plane where the size of the focused laser beam was minimised in the fast (i.e., short confocal parameter) direction.

The differences (errors) between the known defocus offset and the autofocus-predicted defocus correction were calculated for each repeat measurement from each initial defocus offset value. Figures 4(a,b) show the mean and standard deviation of the defocus errors for the shorter range and longer range autofocus predictions respectively. As the initial defocus value was decreased, the error in defocus prediction tends to decrease as expected because the gradients of the autofocus metric curves (Figure 2) increase towards the peak. The autofocus can operate in single-shot functionality over a range of ±37.5 µm (the region where the red plot in figure 2 is above the noise level) and can operate as a 2-step process up to ±68 µm. When imaging a 96-well plate, we found the neighbouring well axial variation to be less than 20 µm, so the accuracy within this region is shown in figure 4(b) in finer defocus offset intervals of 2 µm. Within ±20 µm of focus, the autofocus could correct focus to within the depth of field and within ±5 µm of defocus, the accuracy of the defocus was less than 100 nm. Figures 4(a) and (b) show the results for the long range and short range autofocus performance respectively.

**Figure 4.**
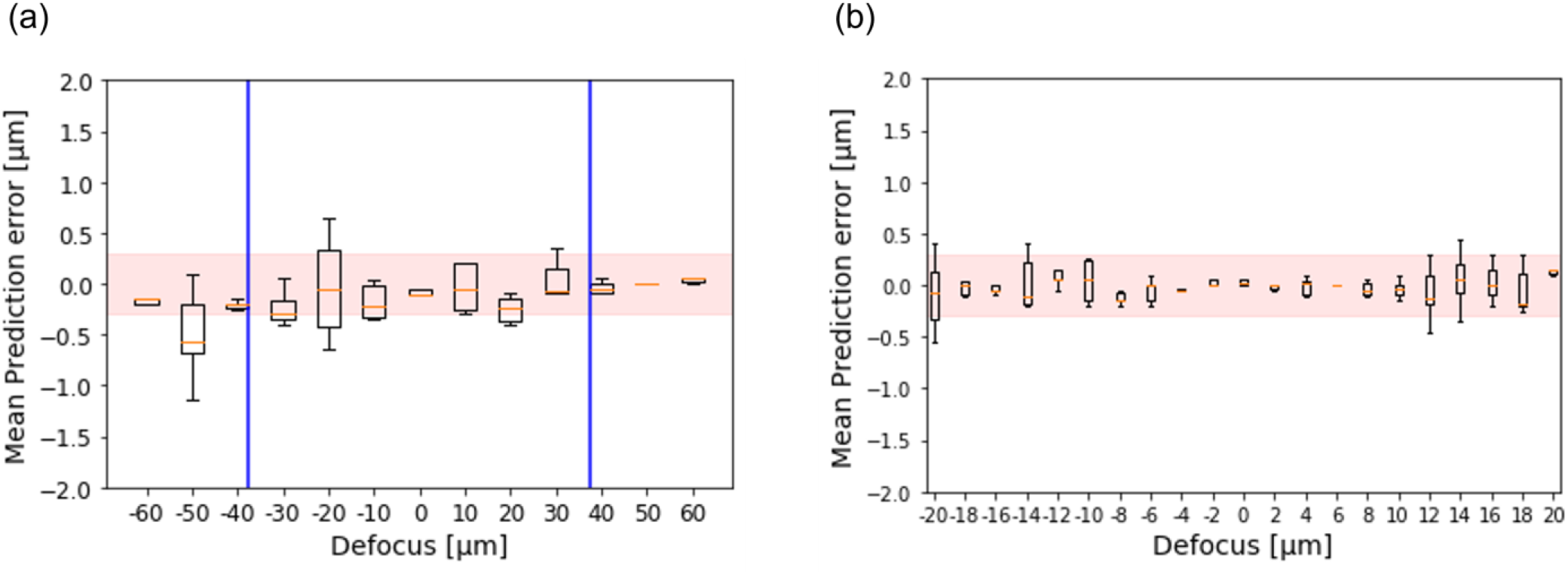
(a,b), System was refocused 10 times from range of known defocus offsets. Errors in predicted defocus are plotted (boxplots with mean error, interquartile range and outliers). Red highlighted region shows depth of field of microscope objective lens. Blue bars indicate the boundaries of the region where autofocus can operate in single-shot

Within the range of the single-shot autofocus capability, it was possible to enable continuous closed-loop focus correction, which is desirable for time-lapse microscopy experiments or when acquiring long image data acquisitions, such as during STORM experiments. If the closed loop autofocus can be implemented independently of the main microscopy imaging, then it will not increase the image data acquisition time. To illustrate the closed-loop autofocus functionality, Figure 5 presents results from a 50,000 frame STORM image data acquisition (over ∼30 minutes at 30 frames/s) that was performed with and without continuous defocus correction to compensate for drift.

**Figure 5.**
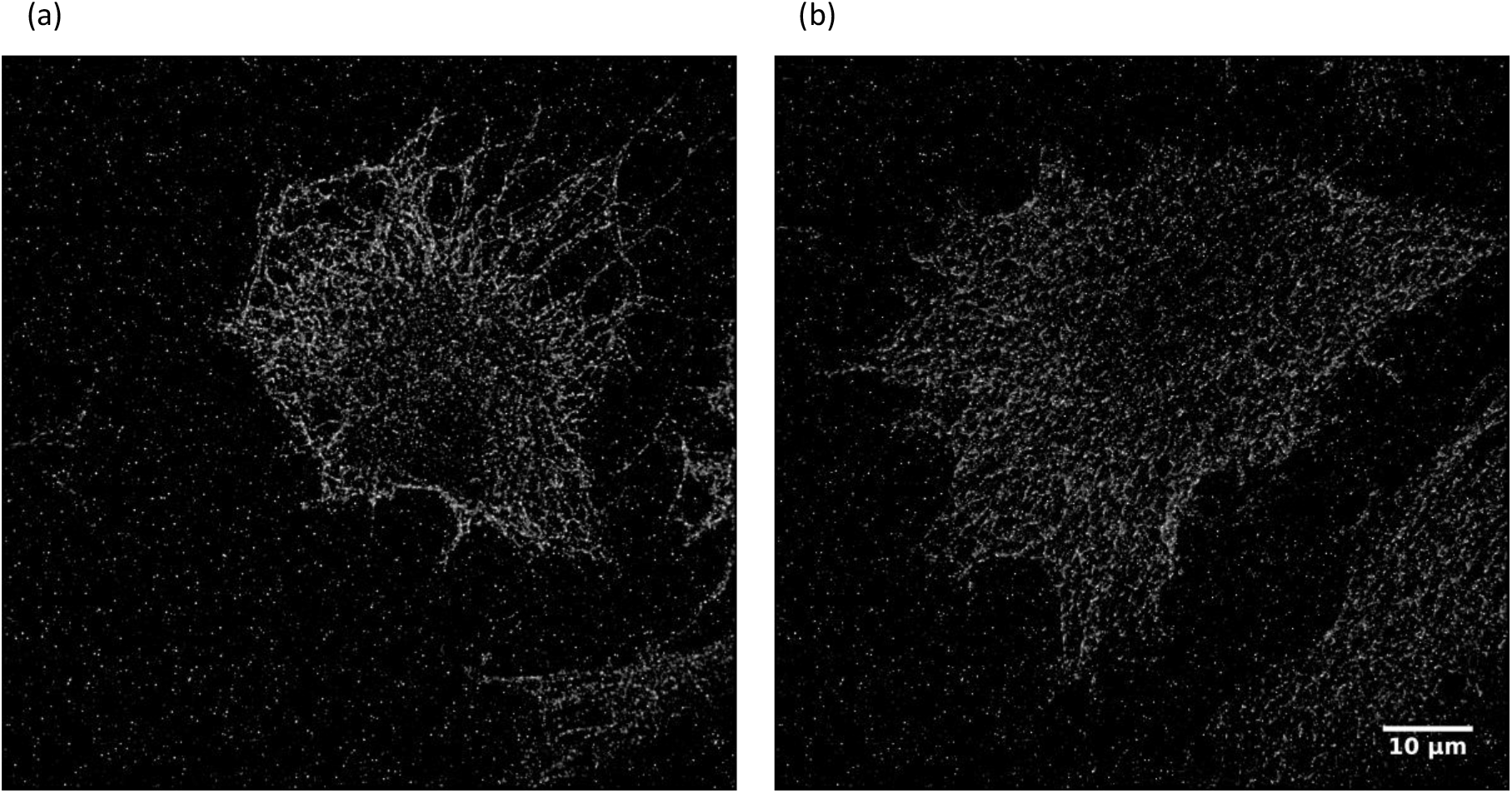
(a) STORM reconstruction of Thyroid cells with the tubulin labelled with ifluor647 acquired on the openFrame using a CellCam kicker camera. Continuous closed loop autofocus correction was used (b) STORM reconstruction of Thyroid cells with the tubulin labelled with ifluor647 acquired on the openFrame microscope using a CellCam kicker camera. Continuous closed loop autofocus correction was not used

The samples were fixed rat thyroid cells (FRTL-5) in which tubulin was labelled with ifluor647 and the easySTORM images were reconstructed using a parallelised implementation of ThunderSTORM running on Imperial’s HPC cluster^13^. Figure 5(a) shows the super-resolved image reconstruction of the easySTORM data acquired when using the continuous closed loop autofocus whereas figure 5(b) shows the reconstruction from the easySTORM acquisition without the closed loop autofocus enabled starting from the microscope in focus. The tubulin filaments are more clearly discernible for the easySTORM acquisition using the closed loop autofocus.

The objective lens stage position was recorded before and after the STORM image data acquisition to quantify the net axial drift. Using the continuous closed loop autofocus, the objective lens stage position changed from 3132.3 µm to 3133.0 µm after the 50000 frames. This axial change of 700 nm exceeds the ±300 nm DOF for this system when using a 635nm excitation source and so the microscope would have drifted out of focus within the STORM image data acquisition without the correction of defocus.

For the case where the autofocus was not used during the acquisition, the objective lens stage position started in focus at 3133.3 µm. The autofocus was then applied after this uncorrected STORM image data acquisition and the in-focus objective stage position was determined to be 3132.9 µm. Thus, the microscope exhibited a system drift of ∼400 nm in the objective lens position during this STORM image data acquisition.

## 4. Conclusions

We have demonstrated a novel robust optical autofocus system that can be added to an existing optical microscope. This addresses the trade-off between autofocus precision and range through the use of cylindrical lenses to adjust the confocal parameter of the autofocus laser beam in orthogonal directions. We have implemented a calibration-based look-up table to determine the displacement of the coverslip from the focal plane of the objective lens and shown that this two-step process works over a range of +/-60 µm with an accuracy of ∼230 nm, which is well within the depth of field of the 1.4 NA objective lens. This approach also provides single-shot autofocusing capability by determining the sign of defocus within the range of the shorter autofocus confocal parameter (±37.5 µm), thereby enabling closed loop “real-time” focus lock. This new approach does not require machine learning but could still benefit from the approach outlined in reference [2], which may extend the operating range as the machine learning can work with lower signal to noise ratio.

## Acknowledgements

Significant components of the instrument reported here were co-designed and fabricated by Simon Johnson and Martin Kehoe in the Optics instrumentation facility of the Physics Department at Imperial College and Tomas Parrado and Jeremy Graham from Cairn Research Ltd contributed to the optomechanical and electronic design of the autofocus module. We gratefully acknowledge funding from the Cancer Research UK ICR/Imperial Convergence Science Centre and Research England GCRF Institutional Award as well as the Imperial College London Impact Acceleration Accounts supported by the Biotechnology and Biological Sciences Research Council (BBSRC EP/R511547/1) and the Engineering and Physical Sciences Research Council (EPSRC EP/R511547/1). We also acknowledge funding from EPSRC (EP/V002910/1) and from Cancer Research UK (A28450, A29368). SK and YA were supported by funding from the Francis Crick Institute, which receives its core funding from Cancer Research UK (FC001003), the UK Medical Research Council (FC001003) and the Wellcome Trust (FC001003). JL acknowledges PhD studentships from EPSRC. For the purpose of open access, the authors have applied a CC BY public copyright licence to this submitted manuscript.

## Notes

### Competing Interest Statement

The authors at Imperial College London are working to develop open source microscopy instruments, partly in collaboration with Cairn Research Ltd, including the openFrame microscope platform and an optical autofocus system for which Cairn have contributed components and technical expertise. Cairn Research Ltd intend to sell components for openFrame microscopes and autofocus systems.

